# Transcriptional regulation of schizophrenia risk gene TCF4 in inhibitory neurons during neurodevelopment

**DOI:** 10.1101/2021.03.04.433878

**Authors:** Yuanyuan Wang, Mingyan Lin

## Abstract

The pathology underlying schizophrenia (SCZ) involves cell type-specific and developmental stage-specific dysregulation of multiple gene regulatory networks dominated by some key transcription factors, such as SCZ risk gene transcription factor 4 (TCF4). Previous studies on the regulatory mechanism of TCF4 use SY5Y as the cellular model, which could not reflect its cell typespecific role in the real world. Using the transcriptional profile of whole brain during development stages and single-cell transcriptome data in the developing human prefrontal cortex, we found that TCF4 was preferentially expressed in the interneuron. Chromatin immunoprecipitation combined with sequencing (ChIP-Seq) in human embryonic stem cells (hESC)-derived interneurons revealed that TCF4 primarily activate transcription of genes associated with cortex development and telencephalon regionalization in a long-range manner. As expected, the downstream targets of TCF4 were distinct in inhibitory neurons and neural stem cells during early neurodevelopment, justifying the importance of our study. Deeper investigation further revealed that TCF4 regulate genes related to neurotransmission distally in interneuron in a c-FOS dependent manner, while TCF4 and TCF3 synergistically regulate genes associated with cell proliferation associated proximally in neural stem cells. Our findings suggested that defects in development of interneuron, for instance as a result of TCF4 abnormality, may break excitation and inhibition balance and contribute significantly to the risk of SCZ.

## Introduction

Schizophrenia is a serious mental illness with complex etiology. It has been increasingly clear that SCZ is influenced by a complex interaction of polygenic genetic and environmental factors^1^, but the pathogenic mechanism is still unclear. Several studies have shown that transcription factor TCF4 is one of the most reproducible susceptibility genes in SCZ-GWAS study^2–4^, showing TCF4 with high consistency in the complex genetic lineage of SCZ. Rare mutations in TCF4 are associated with a range of neurodevelopmental disorders, such as Pitt-Hopkins Syndrome^5^, autism^6^, and bipolar disorder^6^, highlighting a clear link between TCF4 and neurodevelopmental disorders.

By analyzing the transcriptional profiles of the whole brain and the singlecell transcriptome data of the brain during early embryonic development^7,8^, our results showed that TCF4 plays an important role in inhibitory neurons during early neurodevelopment, and TCF4 transcriptional regulation may be cell type specific. However, current researches were mainly based on neural stem cell such as human neuroblastoma cell line SH-SY5Y to investigate the regulation mechanism of TCF4^9,10^. These results reflected the role of TCF4 in neural stem cells, with certain limitations. Therefore, this study will explore the transcriptional regulation of TCF4 in inhibitory neurons in early neurodevelopment and reveal the regulatory mechanism of TCF4 in neurodevelopmental.

## Results

### TCF4 plays a pivotal role in inhibitory neurons in neurodevelopment

To investigate temporal-specific and spatial-specific expression patterns of TCF4 in human brain, we leveraged the transcriptional profile of whole brain during development stages (http://www.brainspan.org/). and analyzed the expression level of TCF4 in each brain development stage. This result showed that the expression level of TCF4 was higher in the early development stage (8 pcw-24 pcw) of four brain regions (Frontal cortex, Sub-cortex, Sensory-motor, Temporal-parietal), and decreased in the postnatal and adult stages (Fig. 1A). Meanwhile, the expression level of TCF4 in the prefrontal cortex was higher than that in the other three brain regions during the early development stage. The observation suggested that TCF4 may play a profound role in the prefrontal cortex during brain development early stage, and the investigation of TCF4 regulatory role should target the early stage of brain development.

**Fig 1.**
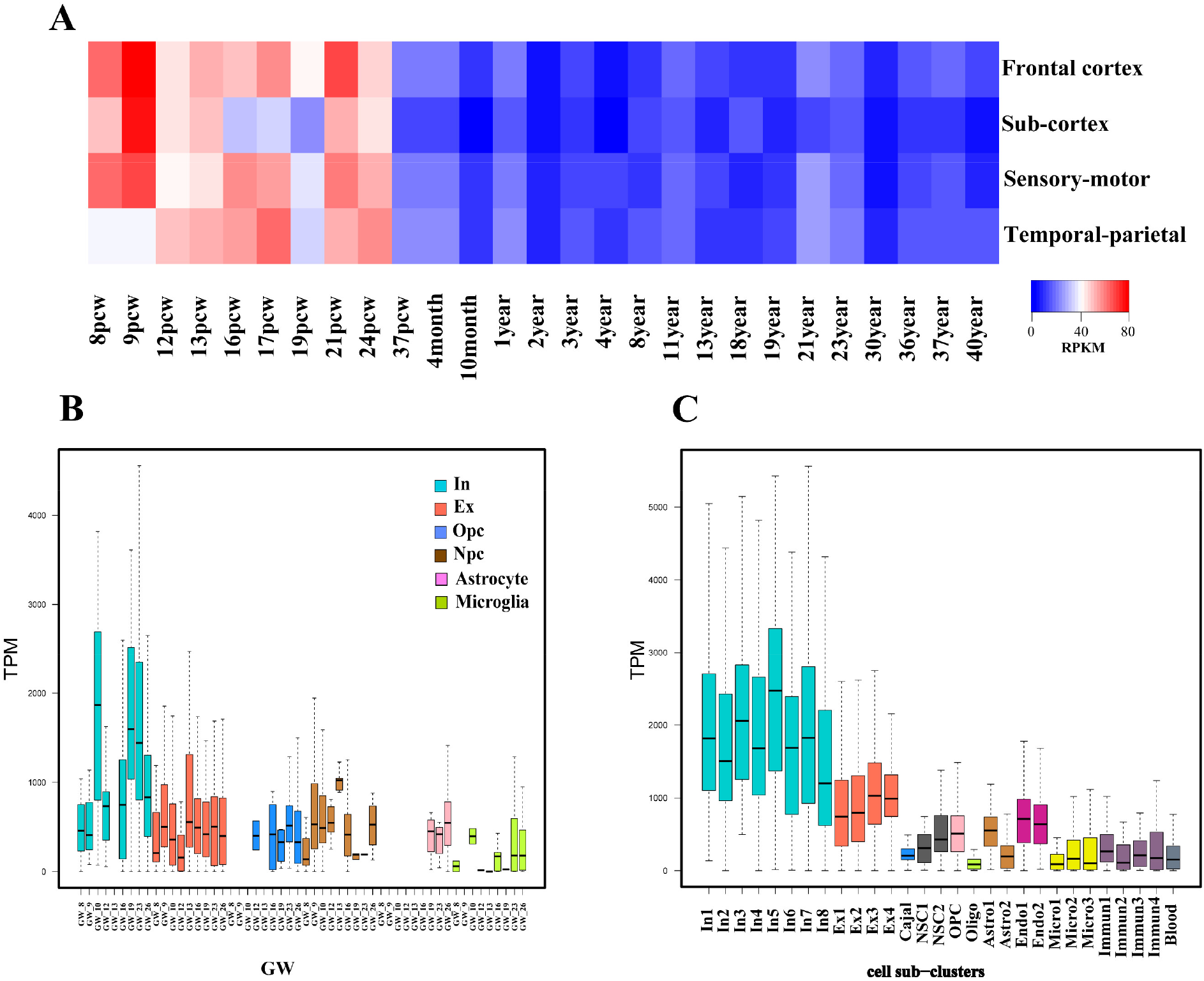
Preferential expression of TCF4 in interneurons. **(A)** Dynamic gene expression profile of TCF4 across different developmental time points and four regions of human brain. The color scale shown on the bottom illustrates the relative expression level of TCF4 across all-time points and brain regions. Red denotes high expression and blue denotes low expression. RPKM: reads per kilobase per million mapped reads. PCW: post-conception week. **(B)** The expression level of TCF4 in 6 cell types of human embryonic prefrontal cortex at gestational weeks (GW)8 to 26. GW: gestational weeks; color: cell types; NPC: neural progenitor cells; Ex: excitatory neurons; In: interneurons; Astrocyte: astrocytes; OPC: oligodendrocyte progenitor cells. TPM: transcripts per kilobase of exon model per Million mapped reads. **(C)** Expression level of TCF4 in 29 cell sub-clusters from the entire human cortex at 22-23 weeks post-conception (22 W and 23 W). Each bar: cell sub-clusters, color: cell types; In: inhibitory neurons; Ex: excitatory neurons; Cajal: Cajal-Retzius cells; NSC: neural stem cell; Opc: oligodendrocyte progenitor cells; Astro: astrocytes; Endo: endothelial cells; Micro: Microglia; immune: immune cells. TPM: transcripts per kilobase of exon model per million mapped reads.

To confirm whether TCF4 expression is cell-type specific, we used singlecell transcriptome data from human embryonic prefrontal cortex at gestational weeks 8 to 26 to analyze the expression level of TCF4 in cell subtypes^7^. The results showed that TCF4 was preferentially expressed in in the interneurons in cerebral cortex cells during early neurodevelopment Compared with other cell types (Fig. 1B). It was speculated that interneuron is the most TCF4-relevant cell type, and TCF4 may act a pivotal part in interneuron. Using another set of high-resolution single-cell transcriptome data from the entire human cortex at 22-23 weeks post-conception (22 W and 23 W)^8^, we verified that TCF4 is preferentially expressed in interneurons (Fig. 1C). These results suggested that TCF4 is a potential key transcription factor in interneurons during early neurodevelopment.

Based on the spatiotemporal- and cell type specific expression of TCF4, it is more appropriate for us to explore the TCF4 regulatory mechanism in interneuron at the early stage of neurodevelopment. Since most inhibitory neurons in human brain originate from the medial ganglionic eminence (MGE) at the early stage of brain development, which migrate to the cerebral cortex and integrate with excitatory neurons to form neural circuit^11–13^. This displayed that the early ventral telencephalon as a model can better fit for the investigation of TCF4 regulatory mechanism.

### Human ES cell-derived interneuron for investigation of TCF4 regulation mechanism

To investigate the regulation mechanism of TCF4 in ventral inhibitory neurons of the early stage of brain development, we followed the protocol previously described in ref^14^. for differentiation of forebrain interneurons from human embryonic stem cells. We performed ventral induce differentiation using hESC H9 cell line to obtain the early interneuron at 26 days of EB differentiation for the subsequent experimental model (Fig. 2A).

**Fig 2:**
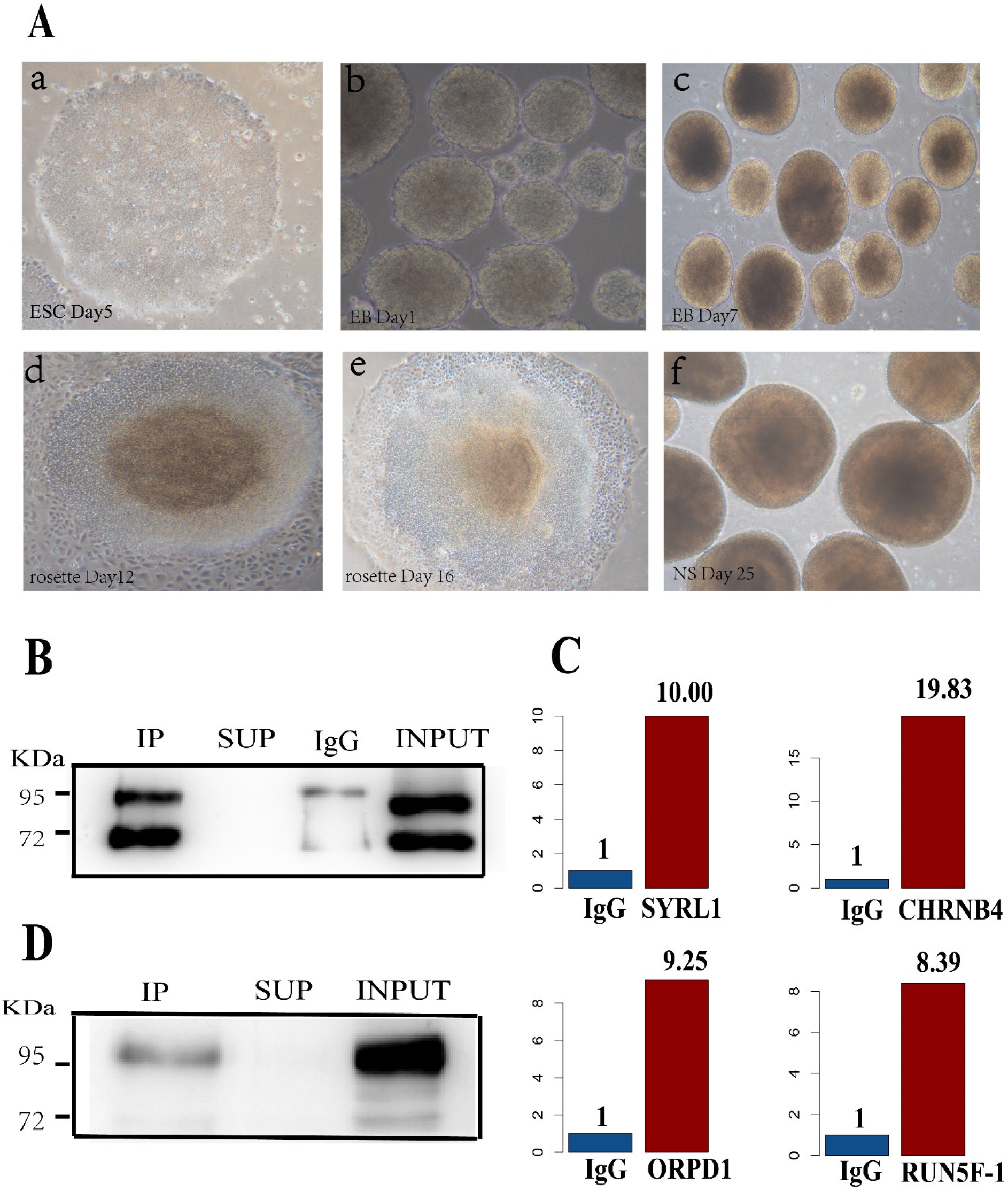
The generation of interneuron in neurodevelopment for TCF4 ChIP assay. (A) Differentiation of forebrain interneurons from human embryonic stem cells (H9 cell lines). **a.** A colony of human ESCs after 5 days in culture showing typical pluripotent morphology. **b.** On day 1, human ESCs were induced to differentiate, cells began to organize into 3D aggregates and continued to grow into spherical EB-like structures. **c.** EBs in suspension culture at the day 7. **d.** The rosettes structure developed from EB on day 12. **e**. The rosettes structure developed from EB on day 16. f. Rosette-containing colonies formed neuroepithelial spheres at day 25. **(B)** Anti-TCF4 antibody efficiently detect TCF4-A (95 KDa) and TCF4-B (71 KDa) from total protein extracts in SH-SY5Y cell line. IP: IP line protein from cell lysis buffer extracts following immunoprecipitation. SUP: The supernatant after extraction of Protein A/G Bead-Antibody/Chromatin Complex. IgG: The negative control Mouse IgG. INPUT: no-antibody ChIP as input. **(C)** The fold enrichment presents ChIP results relative to the negative control IgG. The negative control IgG is given a value of ‘1’and everything else will then be a fold change of IgG. **(D)** Anti-TCF4 antibody efficiently detect TCF4-A (95 KDa) from total protein extracts in interneuron. IP: IP line protein from cell lysis buffer extracts following immunoprecipitation. SUP: The supernatant after extraction of Protein A/G Bead-Antibody/Chromatin Complex. IgG: The negative control Mouse IgG. INPUT: no-antibody ChIP as input.

During the interneuron differentiation procedure, we first generated the embryoid bodies (EB) from the hESC clones with good growth after hESC clones maintain for 5 days (Fig 2A. a). For continuous culture for 7 days, EB in the neural induction medium began to showing spherical structure and starting ectodermal differentiation (Fig 2A. b, c). After 10 days of EB generation, the neural rosettes structure that contains neuroepithelial cells gradually emerged (Fig 2A. d). After 16 days of EB generation, the rosettes-containing colonies are grown in suspension to form neuroepithelial spheres (Fig 2A. e). During the subsequent continuous culture, the neuroepithelial cells begin to differentiate into MGE precursors. and the MGE precursor cells gradually differentiated into GABA interneurons^14^.

Since this study will conduct a comparative investigation on the regulatory mechanism of TCF4 in interneurons and SH-SY5Y cell lines^9^ using ChIP-Seq, we first tested the immunoprecipitation efficiency of the anti-TCF4 antibody in SH-SY5Y cell line in this study and tested whether different batches of antibodies in this study and the previous SH-SY5Y study will cause differences in results. We tested the specificity and ability of anti-TCF4 antibody in SH-SY5Y cells following the ENCODE guidelines^15^ (Fig 2B). Besides, the anti-TCF4 antibody could successfully detect the two major TCF4 isoforms, TCF4-A (95 KDa) and TCF4-B (71 KDa), in SH-SY5Y (Fig 2B).

To confirm the consistency between the enriched Tcf4 binding site DNA fragment and the published SH-SY5Y results, we combined with the binding sites of TCF4 in SH-SY5Y ChIP-Seq that were reported by *Forrest^9^,* and carried out qPCR on the binding sites DNA fragment of TCF4 ChIP-Seq in SH-SY5Y for consistency validation. The candidate TCF4 target genes in SH-SY5Y included CHRNB4, RUN5F-1, OPRD1, and SYPL1. The DNA obtained by ChIP assay with IgG antibody was used as control. The results showed that the genes SYRL1, CHRNB4, ORPD1, and RUN5F-L were significantly enriched in our ChIP assay (Fig. 2 C), which showed that our antibody could successfully enrich the TCF4 binding sites DNA fragment in the previous study in SH-SY5Y cell line. The above results showed that the DNA fragments obtained from ChIP of anti-TCF4 antibody in this study were consistent with the previous results of SH-SY5Y, and would not produce the specific results between different TCF4 antibodies, which ruled out the possibility of differences caused by different antibodies in subsequent comparative analysis.

We tested the specificity and sensitivity of anti-TCF4 antibody in interneurons, and results showed that anti-TCF4 antibody could specifically detect TCF4-A (95 KDa) isoform (Fig. 2 D). This observation showed that TCF4-A was expressed in inhibitory neurons at the early stage of neural development, while both TCF4-A and TCF4-B were expressed in SY5Y. These results indicated that TCF4-A regulates genes in interneuron, whereas TCF4-A and TCF4-B play a role in SY5Y, suggesting that the regulation of TCF4 is cell specific.

### TCF4 target sites identified in interneurons during neurodevelopment

To identify whole genome-wide TCF4 binding sites in interneurons of early neurodevelopment, we ChIP-seq to map TCF4-binding sites in hESC H9-derived interneurons. In interneurons, we identified 5196 high confidence TCF4 peaks. We further screened the peaks contain ‘CANNTG’ classical motif as high credibility TCF4 binding sites for subsequent analysis, and obtained 3460 high confidence TCF4 binding sites. We annotated binding sites with genomic regions using GREAT^16^ and found that most binding sites were located in the distal genomic regions of 50 kb-500 kb away from the transcription start site (Fig 3A, 3B). TCF4 binding sites were enriched at distal genomic regions versus promoter, this suggested that TCF4 may play a long-range regulatory role on target genes in interneurons. We performed a motif enrichment analysis of TCF4 binding sites, and the most enriched *denovo* motif was consistent with the classical motif of the bHLH transcription factor families (Fig 3C).

**Fig 3.**
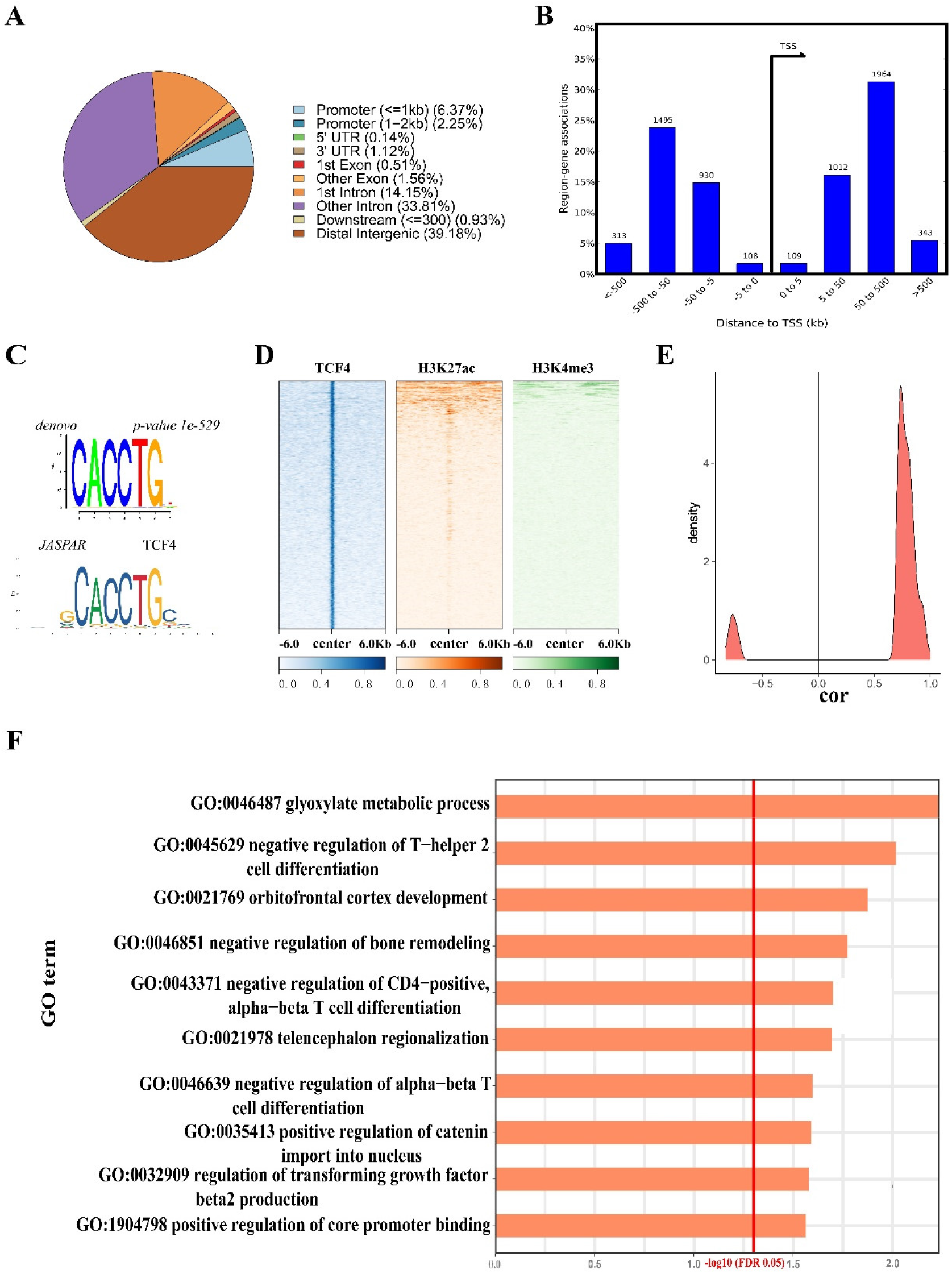
ChIP-seq analysis of TCF4 in interneurons. **(A)** Pie chart of genomic region annotation of TCF4 binding sites. **(B)** Location of TCF4 binding sites relative to transcription start site. **(C)** TCF4 binding sites de novo motif and known motif.TCF4 Known motif was derived from the Jaspar database. **(D)** Heatmap shows read intensity of 50bp in the range from −6 kb to +6 kb centering on TCF4 binding sites from interneurons in TCF4, H3K27AC, and H3K4ME3 ChIP-Seq data. Center: TCF4 genome-wide binding site region. **(E)** Correlation coefficient distribution of TCF4 and TCF4 regulation target gene expression levels in inhibitory neurons in the early neural development of Pearson correlation analysis (p-value < 0.05). The target gene with a correlation p-value less than 0.05 and correlation coefficient greater than 0 was positively correlated with TCF4 expression, while the target gene with a p-value less than 0.05 and correlation coefficient less than 0 was negatively correlated with TCF4 expression. **(F)** Top 10 ranked significant GO terms for TCF4 target genes in interneurons.

### TCF4 activates gene transcription in a long-range manner

To further confirm the long-range regulation of TCF4, we integrated the ChIP-Seq data of active histone modification (H3K27ac and H3K4me3) in fetal brain at 12 weeks of embryonic development to investigate the association between TCF4 binding sites and brain specific active histone modification sites in early brain development^17^. *Fisher’s exact* test enrichment analysis showed that TCF4 binding site significantly enriched the enhancer marker H3K27ac *(p-value* 2.2e-16*, OR=*116.67) of the early fetal forebrain, while the promoter marker H3K4me3 of did not enriched in TCF4 binding site *(p-value* 1) (Fig. 3D). This result suggested that the TCF4 binding sites in interneurons coexist with enhancers in the neurodevelopment stage, which was consistent with the conclusion of genomic region annotation of TCF4 binding sites. The above results emphasized that TCF4 may regulate the transcription of TCF4 target genes in inhibitory neurons by binding to enhancers. To infer target genes regulated by TCF4, we applied GREAT to annotate TCF4 binding sites^16^. We identified 3572 candidate target genes of TCF4, suggesting that TCF4 has a widespread and complex regulatory role.

To determine whether TCF4 can activate or inhibit the transcription of target genes, we carried out pearson correlation analysis on the expression level of TCF4 and 3572 target genes in interneurons in a single-cell transcriptome data of the prefrontal cortex during early brain development^7^. We defined the target gene with a correlation p-value less than 0.05 and a correlation coefficient greater than 0 as the positive correlation gene with TCF4 expression, while the target gene with a p-value less than 0.05 and a correlation coefficient less than 0 as a negative correlation gene with TCF4 expression. Correlation analysis showed that 126 target genes were positively correlated with the level of TCF4 expression, and only 14 target genes were negatively correlated with the level of TCF4 expression (Fig. 3E). The results indicated that TCF4 mainly activated transcription of target genes in interneurons during neurodevelopment.

To elucidate potential biological functions of genes regulated by TCF4, we performed gene ontology enrichment analyses on TCF4 targets. The results of enrichment analysis showed that TCF4 target genes were enriched for functions related to early neural development, such as “cortex development” and “telencephalon regionalization” (Fig. 3F), suggesting TCF4 acts a pivotal part in the cortex development and telencephalon regionalization at neurodevelopment of the early stage.

### The transcriptional regulation of TCF4 was distinct in inhibitory neurons and neural stem cells

Our results showed that the most-relevant cell type of TCF4 is inhibitory neurons. To present the most relevant regulatory mechanism of TCF4 in the early neurodevelopment, we compared the TCF4 ChIP-Seq results in SH-SY5Y reported by *Forrest* with our results^9^.

Firstly, a total of 3752 TCF4 target genes were obtained from interneurons and 5436 TCF4 target genes were obtained from SH-SY5Y. However, there were only 1252 overlapping target genes between interneurons and SH-SY5Y. The target genes of TCF4 were significantly distinct in the inhibitory neurons and SH-SY5Y, and the regulatory genes of TCF4 were different between the two cell types (Fig. 4A). Secondly, TCF4 binding sites in interneurons were mainly located in the distal enhancers of the gene, with a low proportion in the promoters. In SH-SY5Y, the proportion of DNA binding sites in promoter regions is about twice that in interneurons, indicating that TCF4 regulates gene transcription through enhancers in the inhibitory neurons, and regulates gene transcription through promoters in SH-SY5Y (Fig. 4B). Thirdly, we extended 50 bp upstream and downstream of interneurons and SH-SY5Y TCF4 binding sites to identify TCF4 co-factors using Siomics ^18^. Siomics co-factor enrichment analysis showed that transcription factor FOS and AP1 composed of FOS and JUN were significantly enriched in interneurons, and transcription factor TCF3 and MYF were significantly enriched in SH-SY5Y (Fig. 4C, D), suggesting that FOS may be a major cofactor of TCF4 in interneurons. Transcription factor c-fos and c-jun were involved in the regulation of processes in human and higher animals, including those related to neuronal plasticity and immune response. Animal studies have shown that decreased c-fos gene expression in mice directly leads to impaired learning, memory, and synaptic transmission^19^. FOS *rs*1063169, FOS *rs*7101 and JUN *rs*11688 polymorphisms are associated with schizophrenia^20^. Therefore, it is speculated that the synergistic action of c-FOS and TCF4 plays a core role in interneurons. Finally, the sharing TCF4 target genes of interneurons and SH-SY5Y, the specific TCF4 target genes of interneuron, and the specific TCF4 target genes of SH-SY5Y were analyzed by Reactome for pathway enrichment analysis to shows the functions involved in the regulatory mechanism of TCF4 in interneurons and SH-SY5Y^21^. The target genes shared by interneurons and SH-SY5Y share were mainly involved in the proliferation and differentiation of neural stem cells (Fig 4E). TCF4 target genes of interneurons specific enriched for functions related to neurodevelopment, inhibitory neuronal receptors and channel regulation, sodium/calcium ion exchange, neurotransmitter receptors, and postsynaptic signaling pathways (Fig 4F). The TCF4 target genes of SH-SY5Y specific converged on the cell growth pathways such as cell proliferation and cell adhesion (Fig 4G).

**Fig 4.**
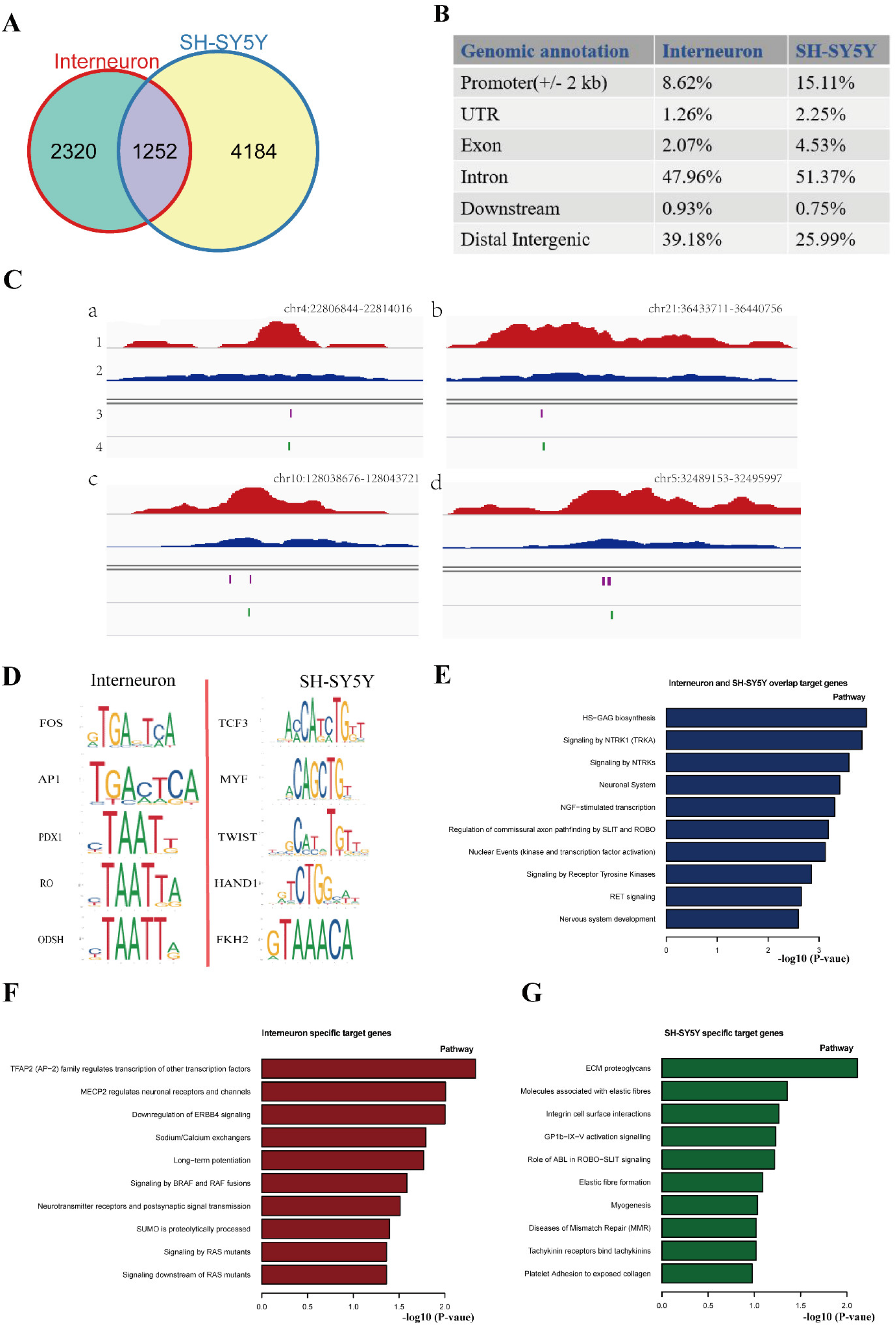
Comparison of TCF4 regulatory role between interneuron and SH-SY5Y. (A) Venn diagram showing the intersection of TCF4 target genes in interneuron with those in SH-SY5Y. The green region represents the TCF4 target genes count of interneuron specific, the yellow region represents the TCF4 target genes count of SH-SY5Y specific, and the purple region is the overlap target genes of interneuron and SH-SY5Y. (B) The genomic region annotation of the TCF4 binding sites in interneuron(left) and SH-SY5Y(right). (C)Integrative Genomics Viewer (IGV) screenshot of TCF4 binding sites and co-factor FOS locus on human chromosome in interneurons. Line 1 within IGV screenshot shows the IP signal. Line 2 within IGV screenshot shows the background signal of available chromatin for IP. Line 3 within IGV screenshot shows TCF4 binding sites locus. Line 4 within IGV screenshot shows TCF4 co-factor FOS binding sites locus. (D) Candidate co-factors of TCF4 in interneuron(left) and SH-SY5Y(right). (E) Top 10 ranked pathways for target genes that overlap between interneuron and SH-SY5Y. X-axis represents -log10(P-value). (F) Top 10 ranked pathways for interneuron specific target genes. X-axis represents -log10(P-value). (G) Top 10 ranked pathways for SH-SY5Y specific target genes. X-axis represents -log10(P-value).

In summary, the regulatory role of TCF4 in inhibitory neurons reflected the more objective regulation mechanism of TCF4 in the early stage of neurodevelopment.

## Discussion

The largest transcriptional analysis of mental diseases has found that many susceptibility genes of neuropsychiatric were specifically enriched in clusters of excitatory neurons in the fetal, which were the key cell types most associated with neuropsychiatric^22^. Our results hypothesized that inhibitory neurons were also key cell types associated with the pathogenesis of psychiatric diseases, and they participated in the occurrence of SCZ together with excitatory neurons.

Genetic studies have found that the effects of SCZ risk variants converge in inhibitory neurons and lead to alterations in the ratio of excitatory to inhibitory cortical activity (E/I imbalance)^23^. Meanwhile, several studies reported the dysregulation of neuropsychiatric risk genes ERBB4^24^, CNTNAP2^25^, NRG1^26^, TSC1^27^,UBE3A^28^, CNTNAP4^29^, and DISC1^30^, which would disturb the function of inhibitory neurons and then lead to E/I imbalance. Intriguingly, these genes are the target genes of TCF4 in inhibitory neurons. Thus, the disruption of TCF4 may contribute to E/I imbalance. Besides, the accumulation of variations of numerous disease risk genes is responsible for the abnormal alteration in excitatory neurons, while in inhibitory neurons, the abnormalities may be caused by the dysregulation of key transcription factors. This difference makes it easier to find widespread intervention targets in inhibitory neurons.

In summary, our study gives valuable insights into the transcriptional regulation of TCF4 in neurodevelopment and implies the link between dysregulation of TCF4 in inhibitory neurons and the pathogenesis of SCZ.

## Author contributions

The study was designed by M.L.. Y.W. performed the experiments. M.L. and Y.W. did the bioinformatic analyses. Y.W., and M.L. wrote the manuscript.

## Acknowledgements

We are grateful to all laboratory members for comments on the manuscript.

